# HIV-1 Capsid Rapidly Induces Long Lived CPSF6 Puncta in Non-dividing Cells but Similar Puncta Already Exist in Uninfected T-cells

**DOI:** 10.1101/2024.03.29.587340

**Authors:** Anabel Guedán, Megan Burley, Eve R. Caroe, Kate N. Bishop

## Abstract

The HIV-1 capsid (CA) protein forms the outer shell of the viral core that is released into the cytoplasm upon infection. CA binds various cellular proteins, including CPSF6, that directs HIV-1 integration into speckle associated domains in host chromatin. Upon HIV-1 infection, CPSF6 forms puncta in the nucleus. Here, we characterised these CPSF6 puncta further, in HeLa cells, T-cells and macrophages and confirm that integration and reverse transcription are not required for puncta formation. Indeed, we found that puncta formed very rapidly after infection, correlating with the time that CA entered the nucleus. In aphidicolin-treated HeLa cells and macrophages, puncta were detected for the length of the experiment, suggesting that puncta are only lost upon cell division. CA still co-localised with CPSF6 puncta at the latest time points, considerably after the peak of reverse transcription and integration. Intriguingly, the number of puncta induced in macrophages did not correlate with the MOI, or the total number of nuclear speckles present in each cell suggesting that CA/CPSF6 is only directed to a few nuclear speckles. Furthermore, we found that CPSF6 already co-localised with nuclear speckles in uninfected T-cells suggesting that HIV-1 promotes a natural behaviour of CPSF6.

## Introduction

The HIV-1 capsid (CA) protein is critical for successful infection. Once the viral envelope has fused with the host cell membrane, the viral core is released into the cytoplasm. The core contains the viral RNA, bound by the nucleocapsid (NC) protein, and the viral enzymes, reverse transcriptase (RT) and integrase (IN). The outer layer of the core is formed of a lattice of CA proteins, mainly assembled into hexamers with 12 pentameric CA arrangements [1–3]. This shell must open or disassemble in order to release the viral nucleic acid before integration can occur but the details of this uncoating process, particularly how, when and where it happens, are poorly defined and somewhat controversial [4]. Uncoating is thought to be triggered by reverse transcription [5–9], but the site of reverse transcription has also recently been re-evaluated and it is now thought that reverse transcription completes in the nucleus [10–12]. Importantly, studies with unstable or hyper-stable CA mutants have shown that viral infectivity depends upon having a core with optimal stability and flexibility [13–15].

HIV-1 can infect non-dividing cells by passing through nuclear pores and the CA protein has been identified as the driver for this nuclear entry [16,17]. This is partly mediated by direct interactions that CA makes with nucleoporins [18–22], however other cellular proteins, including transportins and cleavage and polyadenylation specificity factor 6 (CPSF6, also known as CFIm68) have also been implicated [23–26]. CPSF6 is a nuclear protein involved in alternative polyadenylation of pre-mRNA. Together with CPSF5 (also called CFIm25), it forms the heterotetrameric cleavage factor I mammalian (CFIm) complex [27–29]. CPSF5 can also form heterotetramers with CPSF7 independently of CPSF6, but the functional differences of the two versions of CFIm are unknown. CPSF5 binds RNA and assembles the CFIm complex within a larger Cleavage and Polyadenylation (CPA) complex [27–29]. Following HIV-1 infection, CPSF6 relocates from diffuse nuclear staining to discrete puncta that colocalise with SC35-positive nuclear speckles [12,14,18,30–32]. Nuclear speckles are membrane-less organelles in the nucleus that are involved in mRNA processing and in scaffolding active genome architecture [33]. CPSF6 is suggested to be important to enable HIV-1 cores to pass from the nuclear pore to the nuclear interior [18] and to direct HIV to integrate into active euchromatin in speckle-associated domains (SPADs) [30,31,34,35]. Preventing CA from interacting with CPSF6 results in integration into heterochromatin near the nuclear envelope [30,35,36]. However, the nature of the CPSF6 puncta is only beginning to be defined. Recent work has shown that CPSF6 puncta represent biomolecular condensates that form via liquid-liquid phase separation [37,38] but it is not clear whether CPSF6 needs to accumulate in puncta to facilitate HIV-1 infection. Here, we characterise HIV-1-induced CPSF6 puncta in HeLa cells and primary macrophages and reveal that the puncta form rapidly and continue to be present in cells, associated with CA, for days after reverse transcription and infection occurs. Interestingly, in macrophages, the number of HIV-1-induced CPSF6 puncta present in cells did not correlate with the viral titre or the number of nuclear speckles observed. However, in T-cells, CPSF6 already displayed punctate staining before infection.

## Materials and Methods

### Cells

Adherent 293T and HeLa cell lines were maintained in Dulbecco’s modified Eagle medium (Thermo Fisher). The suspension Jurkat T-cell line was maintained in RPMI-1640 (Thermo Fisher). All cell lines were authenticated and tested mycoplasma-free from Bishop laboratory cell stocks. Media was supplemented with 10% heat-inactivated foetal bovine serum (FBS; Biosera) and 1% Penicillin/Streptomycin (Sigma). Cells were grown in a humidified incubator at 37°C and 5% CO_2_. Peripheral blood mononuclear cells (PBMCs) were isolated from buffy coats from anonymous healthy blood donors (obtained from the NHS Blood & Transplant) by centrifugation using Leucosep tubes following the manufacturer’s instructions (Greiner Bio-One Ltd; 227288). Primary CD4^+^ T-cells and monocytes were purified from PBMCs by positive selection using CD4 (Miltenyi Biotec; 130-045-101) and CD14 (Miltenyi Biotec; 130-097-052) magnetic microbeads, respectively. Primary CD4^+^ T-cells were activated with 100U/μl of interleukin-2 (IL-2; Merck) and phytohemagglutinin-L (PHA-L; Merck) at 2μg/ml for 72h. Primary M2 monocyte-derived macrophages (MDM) were differentiated from monocytes by incubation with macrophage colony-stimulating factor (M-CSF; Miltenyi, 130-096-491) at 25ng/ml for a total of five days. Fresh media containing M-CSF was added at three days post-purification.

In order to arrest HeLa cells at the G1/S boundary of the cell cycle, cells were treated with Aphidicolin (Merck) at 2μg/μl, or DMSO as a control, for 24h before infection. Aphidicolin was maintained in the media throughout the experiment.

### Plasmids and Site-Directed Mutagenesis

The plasmids used to produce HIV-1 viruses, pVSV-G, pCMVΔR8.91 and pCSGW, have been described previously [39,40]. To create Gag-Pol plasmids carrying CA-A14C/E45C, CA-A77V, RT-A114V and IN-D64A, site-directed mutagenesis was performed on pCMVΔR8.91 using the QuickChange II-XL site-directed mutagenesis kit (Agilent) according to manufacturer’s instructions and using the following primers: CA-A14C/*for* 5’- *tttaaagttctaggtgatatgcactgatgtaccatttgcccctg*, CA-A14C/*rev* 5’-*caggggcaaatggtacatcagtgcatatcacctagaactttaaa*; CA-E45C/*for* 5’- *ttgtggggtggctccgcatgataatgctgaaaacatgggtatcactt*, CA-E45C/*rev* 5’- *aagtgatacccatgttttcagcattatcatgcggagccaccccacaa*; CA-A77V/*for* 5’- *gaccatcaatgaggaagttgcagaatgggatagag*, CA-A77V/*rev* 5’- *ctctatcccattctgcaacttcctcattgatggtc*; RT-A114V/*for* 5’- *gtactggatgtgggcgatgtatatttttcagttccctta*, A114V/*rev* 5’- *taagggaactgaaaaatatacatcgcccacatccagtac*; IN-D64A/*for* 5’- *gaatatggcagctagcttgtacacatttag*, IN-D64A/*rev* 5’-*ctaaatgtgtacaagctagctgccatattc*. The introduction of the desired mutations was confirmed by Sanger sequencing (Source Bioscience).

### Single-round Infectious Virus Production

HIV-1-GFP single-round infectious viruses were produced by co-transfecting 293T cells with a 1:1:1 ratio of three plasmids: pVSV-G, pCSGW (GFP-reporter) and pCMVΔ8.91 (or pCMVΔ8.91 mutants). Approximately 16h post-transfection, cells were treated with 10mM sodium butyrate for 8h and VLP-containing supernatants were harvested 24h later. Viral titres were quantified using the Taq-PERT assay to measure viral reverse transcriptase activity [41,42] or, for reverse transcriptase mutants, the Alliance HIV-1 p24 Antigen ELISA from Perkin-Elmer (NEK050B001KT), following the manufacturer’s instructions. GFP expression was detected on a ZOE fluorescent Cell Imager (BIO-RAD).

### Immunofluorescence

Cells were seeded on 13 mm glass coverslips in 24-well plates the day prior to infection. Seeding density was varied depending on the experiment to ensure confluency was not reached before the end of the experiment. Cells were infected with equal RT units/ng p24 of WT or mutant HIV-1-GFP viruses by spinoculation at a low temperature (1600rpm, 16°C for 1.5h) followed by a 30min incubation at to allow synchronized infection. Cells were then washed to remove any unbound virus and incubated at 37°C for the times indicated in the results. Infections resulted in approximately 20% cells infected for T-cells and macrophages and over 80% cells infected for HeLa infections as measured by flow cytometry for GFP in separate experiments. At the indicated times post-infection, cells were washed twice with ice-cold PBS, fixed with PBS supplemented with 4% paraformaldehyde for 5min at room temperature followed by an ice-cold methanol incubation for 5min at -20°C, and washed twice again with ice-cold PBS. After permeabilization with 0.5% saponin (Sigma) in PBS for 30min at room temperature, cells were blocked in 5% donkey serum (DS; Sigma) and 0.5% saponin in PBS for at least 1h at room temperature. Then, cells were incubated with one or more of the following primary antibodies: anti-HIV-1 CA (monoclonal antibody 24-2, a kind gift from Michael Malim), anti-CPSF5 (Proteintech; 66335-1-Ig), anti-CPSF6 (Atlas; HPA039973), anti-CPSF7 (Atlas; HPA041094) and anti-SC35 (Abcam; ab11826), diluted in antibody buffer (1% DS with 0.5% saponin in PBS) for 1h at room temperature. After three washes with PBS, cells were incubated with the following secondary antibodies: donkey anti-rabbit-AF568 (Abcam; ab175692) and donkey anti-mouse-AF647 (Thermo Fisher; A-31571) in antibody buffer for 1h at room temperature. After three washes with PBS, the coverslips were mounted on glass slides (Menzel-Gläser) with ProLong gold antifade mountant with DAPI (Thermo Fisher). Samples were visualised on a SP8 Falcon inverted confocal microscope (Leica) using a 100X oil immersion objective (Leica) or on the IXplore SpinSR super resolution microscope system (Olympus) using an UPLAPO OHR 60X/1.5 oil immersion objective. All image analysis was performed using Fiji.

### Quantitative PCR Analysis to Measure RT Products

Quantitative PCR analysis was conducted as previously described [5,14,43]. Briefly, viruses were treated with 20 units/mL RQ1-DNase (Promega) in 10mM MgCl_2_ for 1h at 37°C before infection. Primary MDM were spinoculated at a low temperature (1600rpm, 16°C for 1.5h) followed by a 30min incubation at to allow synchronized infection. Cells were then washed to remove any unbound virus and incubated at 37°C for the times indicated in the results. Cells were harvested at the indicated times post-infection and total DNA was extracted using the DNeasy Blood & Tissue kit (Qiagen) following the manufacturer’s instructions. The extracted DNA was digested with 1 unit/µL DpnI (Thermo Fisher) for 2.5h at 37°C. qPCR was performed in TaqMan real-time PCR master mix (Thermo Fisher) with 900nM primers and 250 nM probes. The reactions were performed in a 7500 fast real-time PCR system (Applied Biosystems). To calculate DNA copy numbers, standard curves were generated from serial dilutions of pCSGW or a plasmid containing the p2-long terminal repeat (LTR) junction in primary MDM cellular DNA. The following primers and probes were used; minus strand strong stop cDNA products: *for* 5’- *taactagggaacccactgc*, *rev* 5’-*gctagagattttccacactg* and *probe* 5’-*FAM- acacaacagacgggcacacacta-TAMRA*; second strand cDNA products: *for* 5’- *taactagggaacccactgc*, *rev* 5’-*ctgcgtcgagagagctcctctggtt* and *probe* 5’-*FAM- acacaacagacgggcacacacta-TAMRA*; 2-LTR junction: *for* 5’-*gtgtgtgcccgtctgttg*, *rev* 5’- *cagtacaagcaaaaagcagatc* and *probe* 5’-*FAM-ggtaactagagatccctcagacc-TAMRA*.

### Statistics

Statistical analyses were carried out using GraphPad Prism 10 software. Differences between conditions was calculated by one-way ANOVA complemented with Dunnet’s *post hoc* test (*, P < 0.05; **, P < 0.01; ***, P < 0.001; ****, P < 0.0001).

## Results

### HIV Infection Re-Localises CPSF6 into Puncta in Dividing and Non-Dividing Cells

In order to investigate CPSF6 function during HIV-1 infection, we first infected a range of cell types with HIV-1-GFP, immunostained for CPSF6 and the nuclear speckle marker SC35 and imaged by confocal microscopy (Figure 1A & 1B, Figure 2A and Figure S1A). The fluorescent signal intensity of each marker was quantified along an arbitrary line through each cell to assess the degree of dispersion and co-localisation (Figure 1C & 1D, Figure 2B and Figure S1B). Figure 1A & 1B shows that CPSF6 (magenta) generally localised throughout the nucleus of HeLa cells and macrophages, with the exception of the nucleoli, in a diffuse pattern in uninfected cells and then reorganised into distinct puncta that co-localised with SC35 (white) upon infection (Figure 1A-D). However, it was noticeable that CPSF6 staining was already less diffuse and more punctate in uninfected Jurkat T-cells compared to the other cell types (Figure 2A and S1A). Moreover, the CPSF6 staining observed in uninfected cells colocalised with SC35 speckles (Figure 2B and S1B). To confirm that this was not an artifact of immortalised T-cell lines, we also stained activated primary CD4+ T-cells for both CPSF6 and SC35 and observed similar puncta in these T-cells (Figure S2). The high background of CPSF6 puncta in uninfected T-cells made it harder to detect puncta caused by infection and there was no increase in the percentage of cells with puncta or the number of puncta per cell following infection (Figure 2C). However, we did notice that CPSF6 staining was often in smaller, bright foci that resembled the HIV-induced relocation of CPSF6 in other cell types after infection (Figure 2 S1, blue arrows). Therefore, HIV-1 infection re-localises CPSF6 to SC35-containing nuclear speckles but in T-cells, CPSF6 already accumulates to some extent at these sites in the absence of infection.

**Figure 1.**
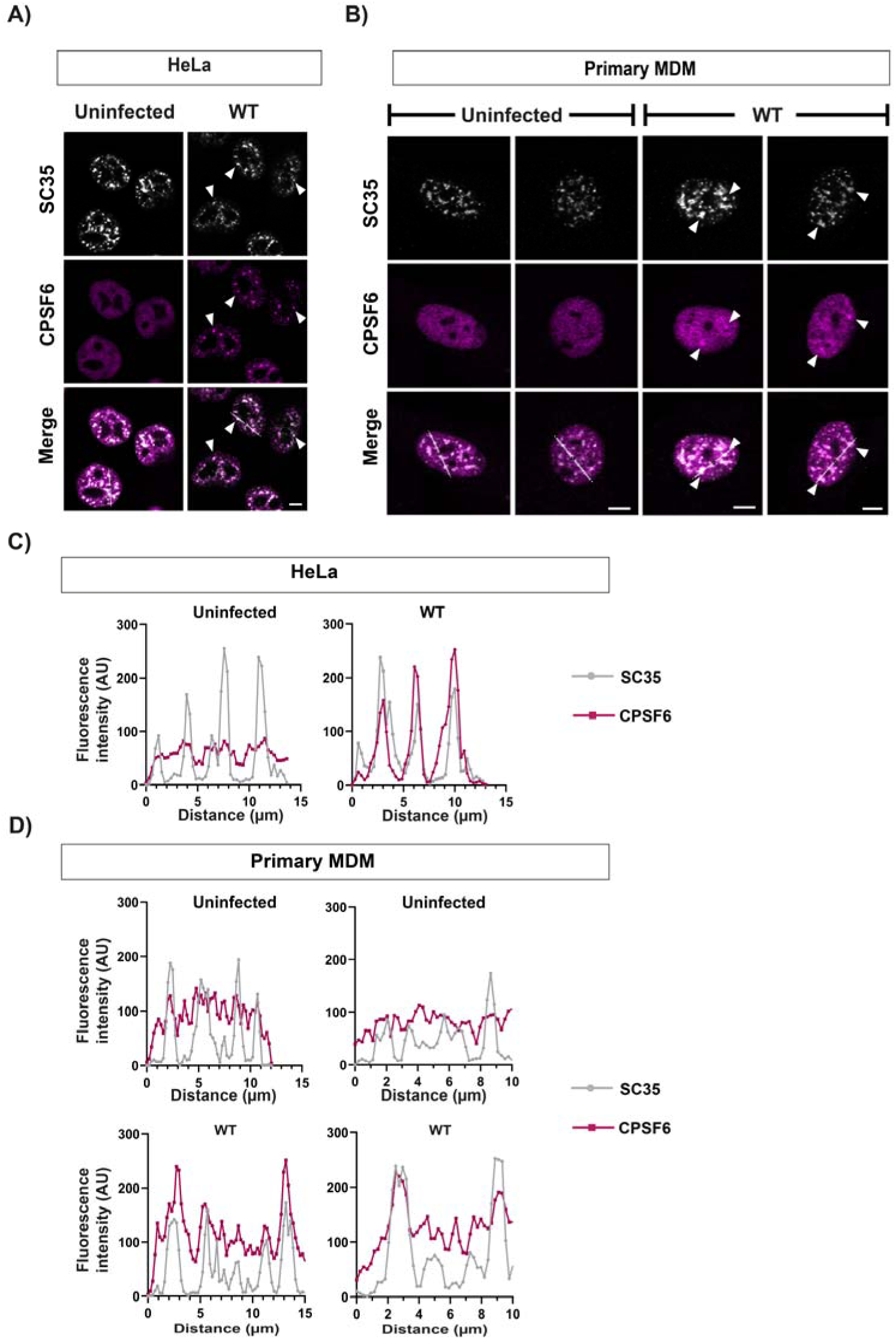
HIV-1 CA induces re-localisation of CPSF6 to nuclear speckles. HeLa cells (A, C) or primary MDMs (B, D) were infected with equal RT units of wild type (WT) HIV-1-GFP and fixed at 16 hours post infection (h.p.i.) or 5 days post infection (d.p.i.), respectively. Uninfected and infected cells were immunostained with primary antibodies against SC35 and CPSF6 followed by species-specific secondary antibodies conjugated to Alexa Fluor fluorophores. (A-B) Representative images of SC35 (white) and CPSF6 (magenta) staining. White arrowheads point to SC35 and CPSF6 co-localisation. Scale bars are 5μm. (C-D) Representative intensity profiles of SC35 and CPSF6 staining in HeLa (C) and primary MDMs (D). The graphs show the fluorescence intensity along the white lines shown in the merged images in (A) and (B), respectively.

**Figure 2.**
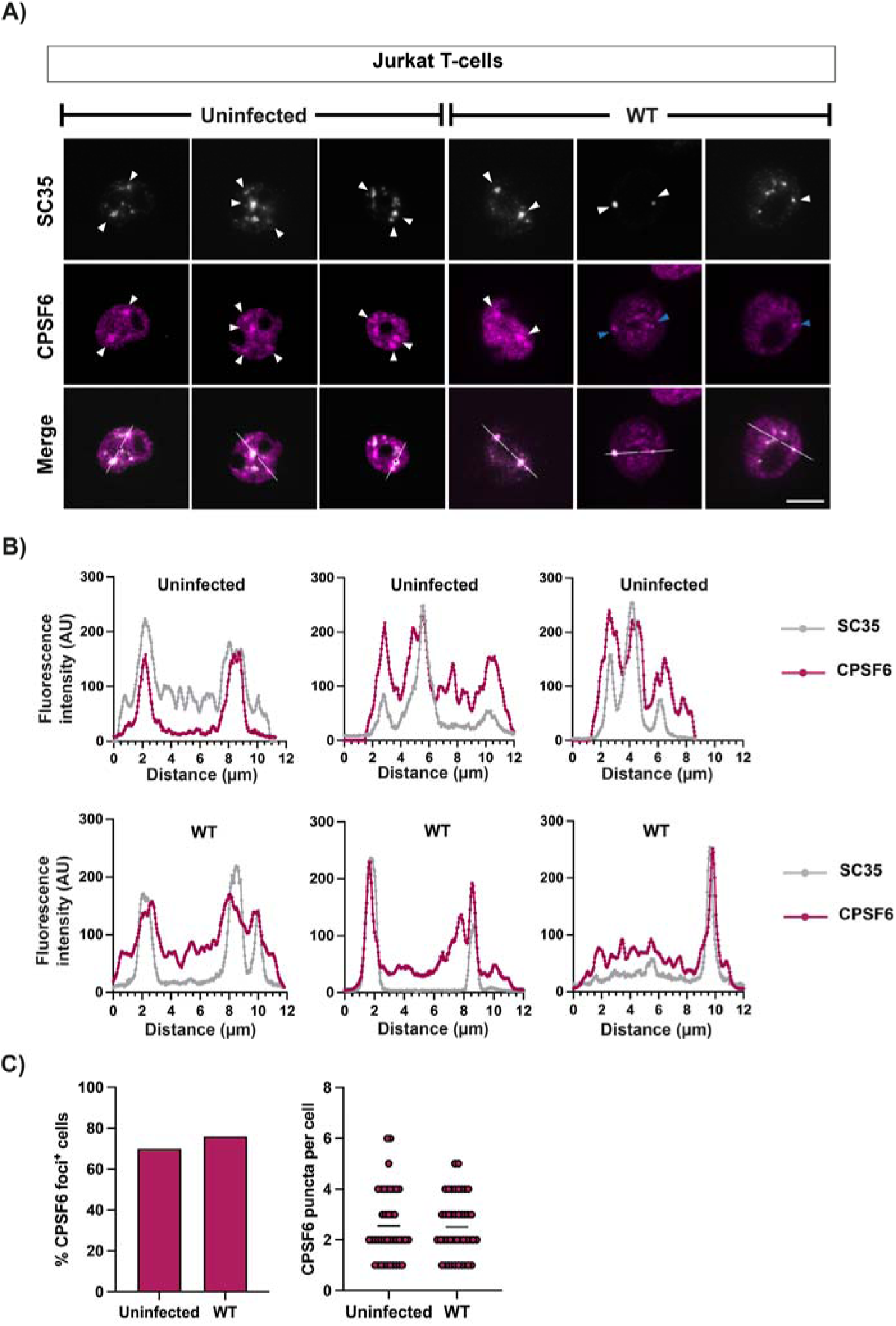
CPSF6 staining is punctate in uninfected Jurkat cells and co-localises with nuclear speckles. Jurkat T-cells were infected with WT HIV-1-GFP and fixed at 24 h.p.i. Uninfected and infected cells were immunostained with primary antibodies against SC35 and CPSF6 followed by species-specific secondary antibodies conjugated to Alexa Fluor flurophores. (A) Representative images of SC35 (white) and CPSF6 (magenta) staining. White arrowheads point to SC35 and CPSF6 co-localisation. Blue arrowheads point to small, bright foci. Scale bars are 5μm. (B) Representative intensity profiles of SC35 and CPSF6 staining in uninfected (top graphs) and infected (bottom graphs) Jurkat T-cells. The graphs show the fluorescence intensity along the white lines shown in the merged images (A). (C) Quantification of the percentage of cells positive for CPSF6 foci (left graph) and the number of CPSF6 puncta per cell (right graph).

### CPSF5 but Not CPSF7 Is Relocalised with CPSF6

CPSF6 is part of a large CPA complex. It binds to CA by virtue of an Phe-Gly (FG) peptide that engages the FG-binding pocket formed between the N-terminal domain (NTD) of one CA monomer and the C-terminal domain (CTD) of the adjacent CA monomer in the CA hexamer [44,45]. So far, it is the only CPA protein known to interact directly with an HIV-1 protein. To determine whether other components of the CPA complex also relocate with CPSF6, we infected HeLa cells and primary MDMs with HIV-1-GFP and immunostained for CPSF6 and its binding partner in the CFIm complex, CPSF5 (Figure 3A & 3B and Figure S3A). CPSF5 showed similar diffuse nuclear staining as CPSF6 in both uninfected HeLa cells and MDMs (Figure 3B and Figure S3A) and re-localised to distinct puncta following infection. There was absolute co-localisation of CPSF5 with CPSF6 in puncta in both cell types (Figure 3B and Figure S3A) indicating that CPSF6 relocates as a complex rather than an individual protein in both cycling and non-cycling cells.

**Figure 3.**
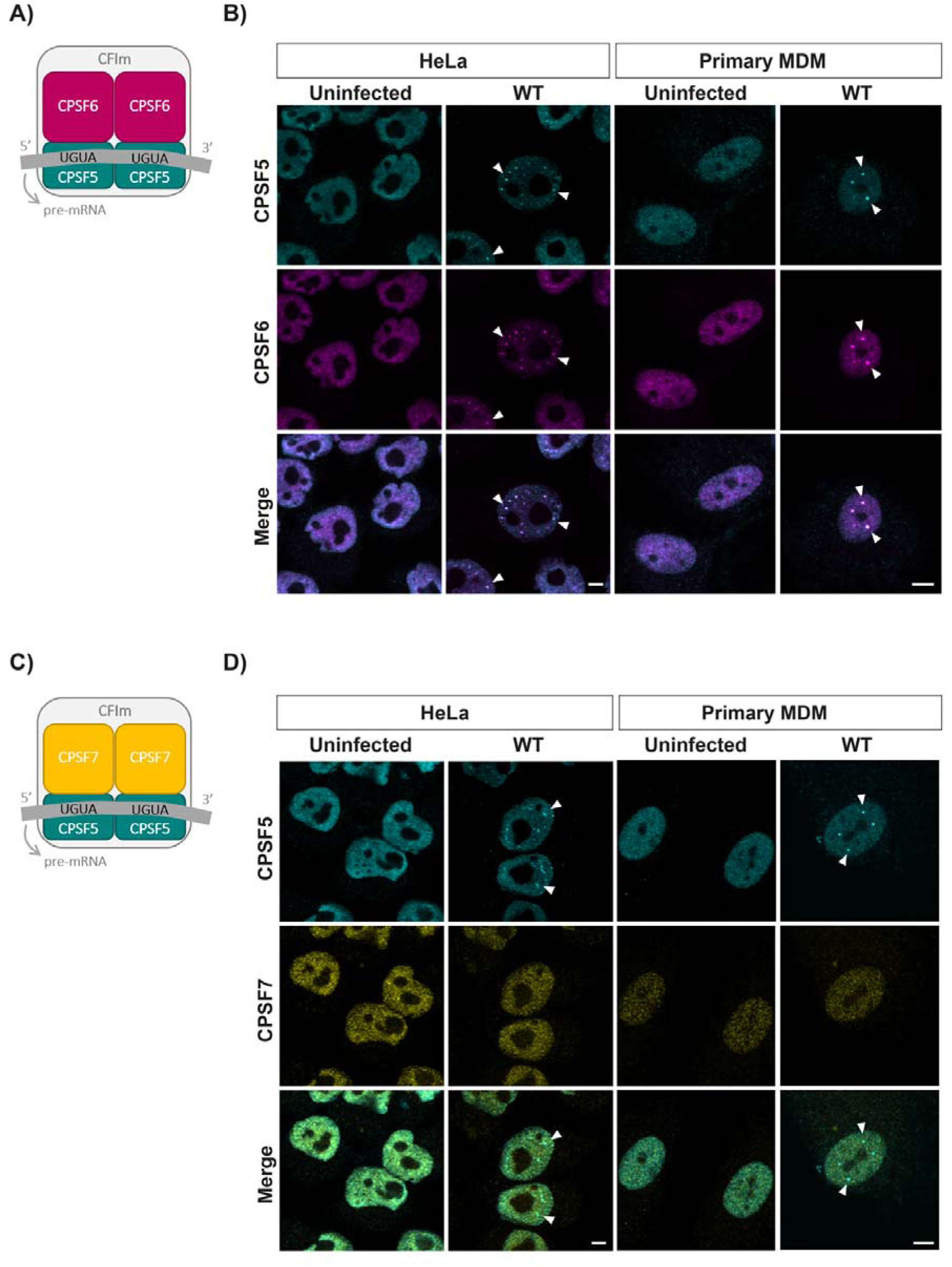
HIV-1 CA induces relocalisation of CPSF5 and CPSF6 but not CPSF7. (A) and (C) Schematic diagrams of the CFIm complexes with CPSF5 and CPSF6 (A), and CPSF5 and CPSF7 (C). (B) and (D) HeLa cells and primary MDMs were synchronously infected with equal RT units of WT HIV-1-GFP and fixed at 16 h.p.i. and 72 h.p.i., respectively. Representative images of uninfected and infected cells immunostained for CPSF5 and CPSF6 (B) or CPSF5 and CPSF7 (D) are shown. White arrowheads point to co-localisation. Scale bars are 5μm.

We next immunostained cells with another CFIm complex protein, CPSF7. Like CPSF6, CPSF7 can form a hetero-tetramer with CPSF5 (Figure 3C). Due to antibody species, we were unable to co-stain for CPSF6 but we could co-stain for CPSF7 and CPSF5 (Figure 3D and Figure S3B). Although CPSF7 had a more grainy staining pattern in both HeLa cells and MDMs, it did not change upon infection with HIV-1, and it did not co-localise with the CPSF5 puncta that formed following infection in either cell type (Figure 3D and Figure S3B). Thus, only the CPSF6-containing CFIm complex rearranges in the presence of HIV-1.

### Reverse Transcription and Integration Are Not Required for Puncta Formation

As the re-organisation of the CFIm complex appeared to be dependent on the direct interaction of CPSF6 with HIV-1 CA, we next tested which steps of HIV-1 replication were required for CPSF6 puncta formation. We infected HeLa cells and MDMs with HIV-1-GFP carrying specific mutations, immunostained cells for CA and CPSF6 and imaged cells by confocal microscopy (Figure 4). Firstly, we introduced the A77V substitution in CA that inhibits CPSF6 binding to the FG-binding pocket of CA [46]. We also introduced the A114V substitution in RT or the D64A substitution in IN that inhibit the catalytic activity of each viral enzyme respectively [47,48]. Finally, we included a mutant with two cysteine substitutions in CA that induce disulphide bond formation across the NTD of CA and stabilises the CA lattice, that we previously showed retarded nuclear import, although it can still bind CPSF6 *in vitro* [14]. Blocking CA binding to CPSF6 (CA-A77V) or stabilising the CA lattice causing cores to get stuck in the nuclear pore (CA-A14C/E45C) prevented puncta formation, even in dividing cells (Figure 4A & 4B), indicating that CA must interact directly with CPSF6 to induce CPSF6 reorganisation. However, inhibiting integration (IN-D64A) had no effect on puncta formation (Figure 4) and CPSF6 puncta even formed in the absence of reverse transcription in both dividing and non-dividing cells (Figure 4), showing that this phenomenon is independent of viral replication and only requires CA to be present in the nucleus.

**Figure 4.**
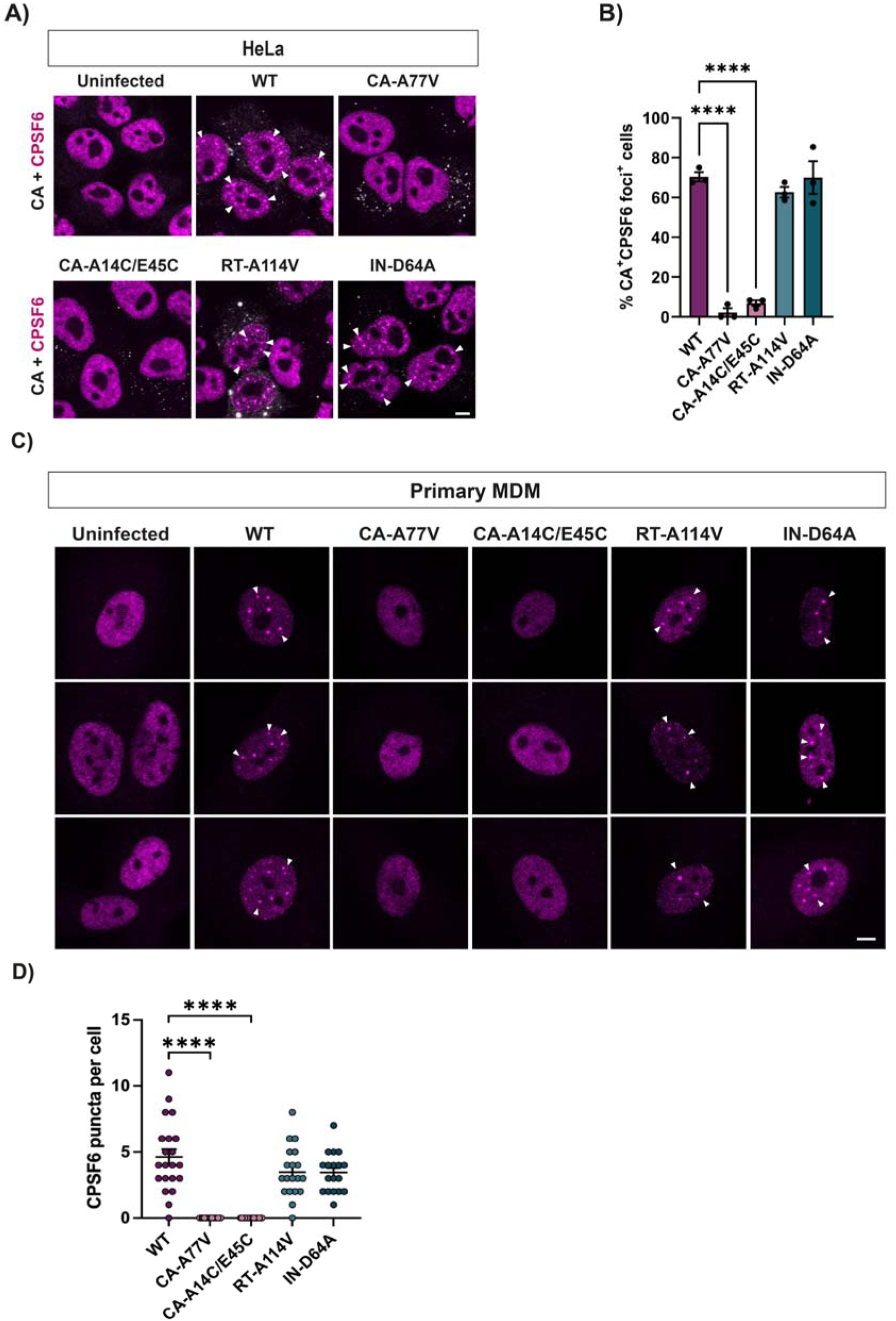
CPSF6 relocalisation is independent of reverse transcription and integration. (A) HeLa cells were infected with equal titres of WT or mutant HIV-1-GFP (CA-A77V, CA-A14C/E45C, RT-A114V and IN-D64A) for 16 h.p.i. Uninfected cells were used as a control. Cells were incubated with primary antibodies against HIV-1 CA (white) and CPSF6 (magenta) followed by species-specific secondary antibodies conjugated to Alexa Fluor fluorophores. Representative merge images are shown. The scale bar is 5μm. (B) Bar chart represents the percentage of CA positive cells that showed CPSF6 redistribution to puncta. Points indicate individual biological repeats (∼150 cells per repeat were analysed for quantification) and bars show the mean ± SEM; **** = p<0.0001, analysed by one-way ANOVA with Dunnet’s multiple comparisons test. (C) Primary MDMs were synchronously infected with equal RT units of WT or mutant VLP (CA-A77V, CA-A14C/E45C, RT-A114V and IN-D64A) for 72 h.p.i. Cells were immunostained for CPSF6. Three representative images are shown per condition. Scale bar is 5μm. (D) Quantification of the number of CPSF6 puncta per cell in infected MDMs.

### CPSF6 Puncta Form Rapidly and Decay Slowly in HeLa Cells

As CPSF6 puncta are induced by CA in the nucleus independently of reverse transcription and we previously showed that CA can be detected in nuclear fractions within the first two hours of infection in HeLa cells [14], we wondered how long it took CPSF6 puncta to form. We therefore synchronously infected HeLa cells with equal amounts of wild type (WT) and RT-A114V HIV-1-GFP and imaged cells for CA and CPSF6 puncta at different times post-infection (Figure 5A and Figure S4). The number of CA positive cells with CPSF6 puncta are plotted for each time point in Figure 5A. Confirming that CPSF6 puncta formed independently of reverse transcription, the kinetics of puncta formation were the same for WT and RT-A114V HIV-1 infections (Figure 5A). Interestingly, the number of puncta increased rapidly with time until 8 hours post-infection (h.p.i.), then plateaued and started declining slowly from 24 h.p.i. (Figure 5A). This correlates well with our published data for the timing of CA arrival at the nucleus: Using immunoblotting for CA in fractionated cell lysates and also proximity ligation assays for nuclear pore proteins and CA, we previously showed that A14C/E45C-CA could be detected in the nuclear fraction of cell lysates as early as 30 minutes post-infection although WT CA was only detected at 2 h.p.i. and peaked at 8 h.p.i. [14]. The timing of CPSF6 puncta formation in HeLa cells also correlated with the timing of reverse transcription in WT HIV-1 infected cells (Figure 5A and [14]).

**Figure 5.**
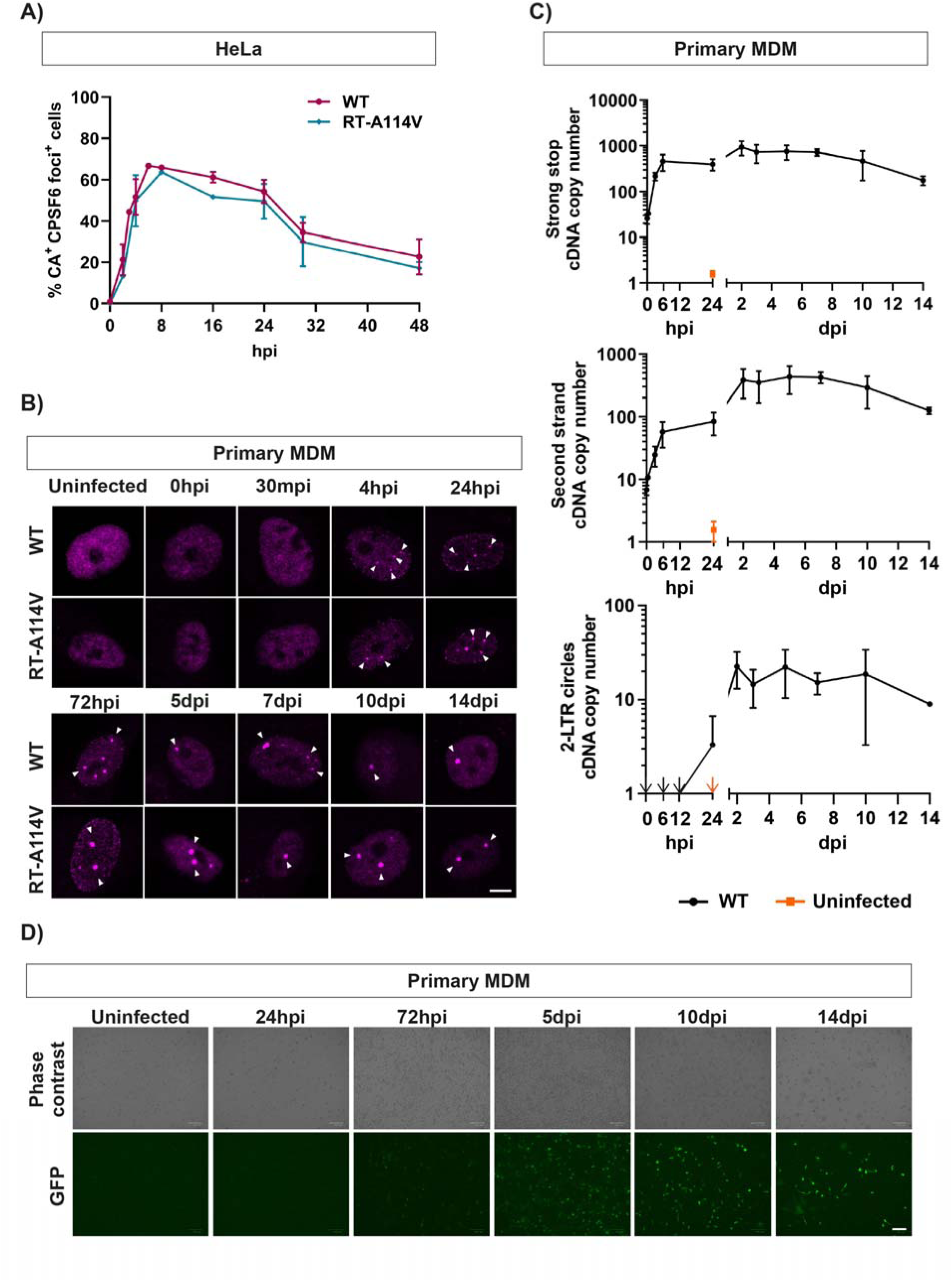
CPSF6 puncta form rapidly and decay slowly in HeLa cells but can exist over two weeks in primary MDM. (A) HeLa cells were synchronously infected with equal ng p24 of WT and RT-A114V HIV-1-GFP, and fixed at 0, 2, 4, 8, 16, 24, 30 and 48 h.p.i. Cells were immunostained for HIV-1 CA and CPSF6. The graph displays the percentage of CA positive cells that showed CPSF6 redistribution to puncta. Data is plotted as mean ± SEM of at least three biological repeats (∼150 cells per repeat were analysed). (B) Primary MDM were synchronously infected with equal ng p24 of WT and RT-A114V HIV-1-GFP, and fixed at 0h, 30min, 4h, 24h, 72h, 5d, 7d, 10d and 14 d.p.i. Cells were immunostained for CPSF6. Representative images from three biological repeats are shown. Scale bar is 5μm. (C, D) Primary MDM were synchronously infected with equal RT units of WT HIV-1-GFP, and harvested for DNA extraction at 0h, 30min, 3h, 6h, 24h, 48h, 72h, 5d, 7d, 10d and 14 d.p.i. (C) Early (minus strand strong stop, top panel) and late (second strand, middle panel) viral cDNA products, and 2-LTR circles (bottom panel) were measured by qPCR. The data is shown as mean ± SEM of three biological repeats. Arrows indicate data points that lie out of range of the y axis. (D) Representative images of GFP-expressing cells (bottom panels) at some of the time points shown in (C). Pictures were taken before harvesting the cells for DNA extraction. The scale bar is 100μm.

### CPSF6 Puncta Form Rapidly and Can Exist for over Two Weeks in Macrophages

We then repeated the time course experiment in primary MDMs. We synchronously infected MDMs with equal amounts of WT HIV-1-GFP and harvested cells at different times post-infection. Replication is known to be slower in these cells, probably partly due to decreased dNTP levels [49], so cells were imaged over a longer time course, over two weeks post infection (Figure 5B). In these infections, CPSF6 puncta (magenta) can clearly be seen as early as 4 h.p.i. The puncta appeared to increase in size/brightness until 72 h.p.i. but, importantly, they remained present for at least two weeks (when the cells started to die), regardless of whether the particles could reverse transcribe or not (Figure 5B). To compare the kinetics of puncta formation to the kinetics of HIV-1 replication in MDMs, we repeated the infections and harvested cells for DNA extraction at different times post-infection (Figure 5C). Cells were imaged for GFP expression from the GFP-reporter gene carried by the virus before harvesting (Figure 5D). Viral cDNA was analysed by qPCR for early (minus strand strong stop) and late (second strand) reverse transcription products and 2-LTR circles (Figure 5C). Strong stop cDNA levels increased rapidly until 6 h.p.i. and then plateaued (Figure 5C, top panel). However, second strand cDNA continued to increase until 48 h.p.i. before levels plateaued (Figure 5C, middle panel). This is expected as reverse transcription can start in viral particles with incorporated dNTPs from producer cells whereas the synthesis of longer products requires dNTPs from target cells [50,51]. In keeping with the delayed reverse transcription in MDMs, 2-LTR circle production that represents completion of reverse transcription also increased until approximately 48 h.p.i. (Figure 5C, bottom panel). Reverse transcription, integration, cellular transcription and translation all need to occur before reporter genes can be detected. Consistent with this, GFP expression was first detected at 72 hours and peaked around 5 days post infection (Figure 5D). Altogether, this shows that CPSF6 puncta form rapidly in MDMs, before reverse transcription peaks but remain in these cells for days after the peak times of integration and subsequent gene expression.

### CPSF6 Puncta Remain until Cells Divide and Are Associated with CA Even at Late Times

CPSF6 puncta formed rapidly in both HeLa cells and MDMs but the puncta stability appeared to differ in the two cell types. In non-dividing MDMs, puncta were present for at least two weeks post infection whereas in dividing HeLa cells the puncta started to disappear after 24 h. This could be because during mitosis, the nucleus is broken down and reformed which may remove any puncta present. We therefore repeated the synchronised infections with HeLa cells that had been treated with aphidicolin to block cell division (Figure 6A). In these cells, the puncta still formed rapidly, with similar kinetics to those seen in untreated HeLa cells, but now the puncta remained for at least 48 h, until the cells started to die from the aphidicolin treatment (Figure 6A). Again, the kinetics were the same between VLPs that could and could not reverse transcribe. Thus, CPSF6 puncta seem very stable as long as the cells do not divide. Interestingly, we noticed that CA was associated with CPSF6 puncta in aphidicolin-treated HeLa cells even at late time points (Figure 6B). When we analysed the puncta in primary MDMs we found that CA was also associated with CPSF6 puncta at late time points in these cells (Figure 6C-D). This suggests that the CA protein stably interacts with CPSF6 puncta, although it is not possible from these experiments to determine how much of the CA lattice was present nor whether there was any viral nucleic acid associated with CA or the CPSF6 puncta.

**Figure 6.**
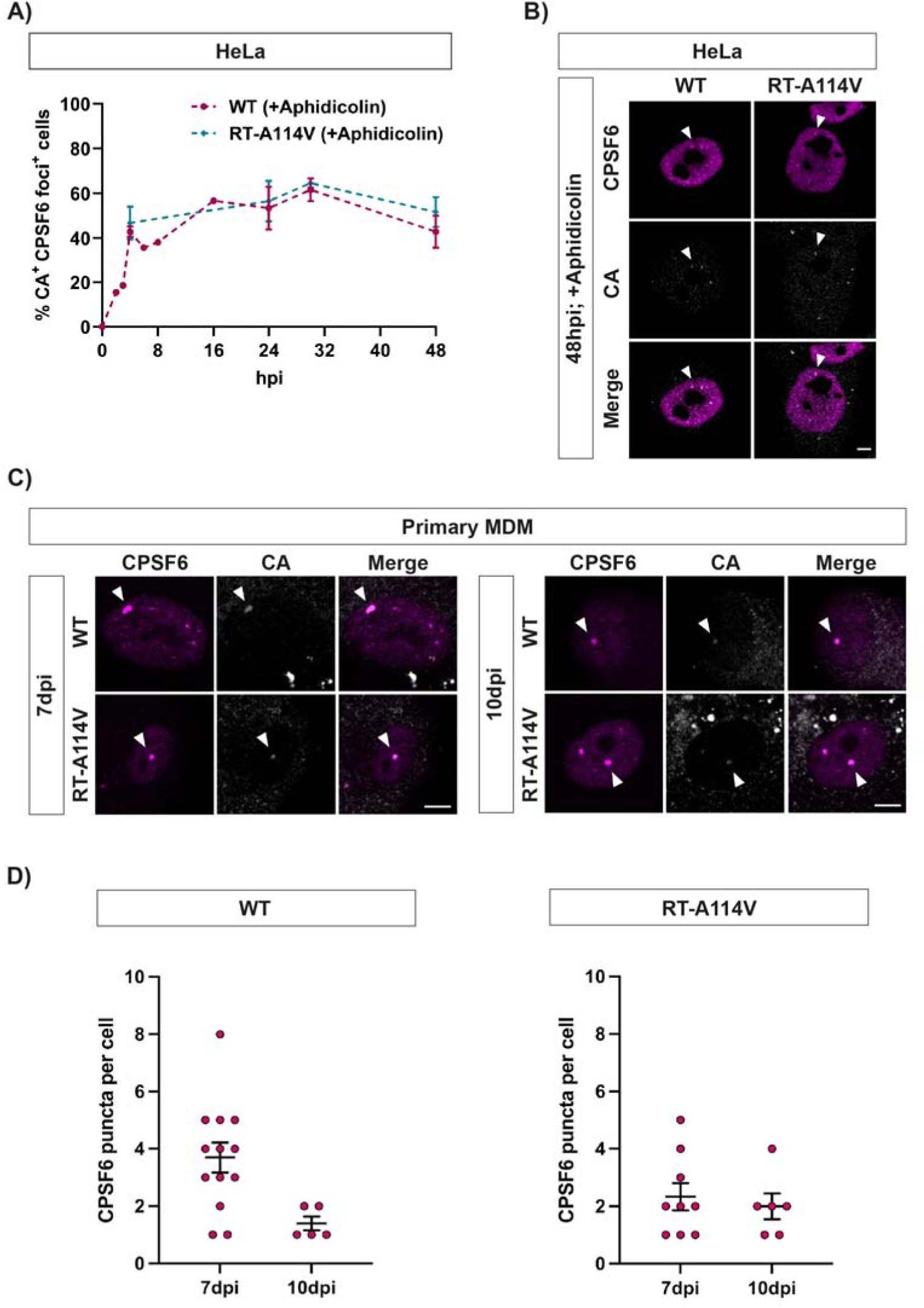
CPSF6 puncta remain in non-dividing cells and are associated with CA even at late times. (A, B) HeLa cells were arrested in S phase by pre-treatment with Aphidicolin (2μg/μL) for 24h prior to a synchronous infection with equal RT units of WT or RT-A114V HIV-1-GFP. Aphidicolin was kept in the media throughout the experiment. Cells were fixed at 0, 4, 24, 30 and 48 h.p.i. and immunostained for HIV-1 CA and CPSF6. (A) The graph displays the percentage of CA positive cells that showed CPSF6 redistribution to puncta. Data is plotted as mean ± SEM of 2-3 biological repeats (∼150 cells per experiment were used for quantification). (B) Representative images of Aphidicolin-treated cells infected with WT or RT-A114V HIV-1-GFP for 48h. (C) Primary MDMs were synchronously infected with equal RT units of WT or RT-A114V HIV-1-GFP and fixed at 7 and 10 d.p.i. Cells were immunostained for HIV-1 CA and CPSF6. Representative images are shown. (B) and (C) White arrowheads point to HIV-1 CA and CPSF6 co-localisation. Scale bar is 5μm. (D) Quantification of the number of CPSF6 puncta per cell in infected MDMs at 7 and 10 d.p.i.

Although CPSF6 appeared to be punctate in uninfected Jurkat T-cells, once the cells had been infected, we noticed an increase in smaller and more focused puncta (Figure 2, blue arrows). Therefore, we infected Jurkat T-cells with HIV-1-GFP and followed the CPSF6 puncta with time (Figure S5). Interestingly, we detected the smaller puncta at the earliest time point we imaged (2 h.p.i.), and they were visible until 24 h.p.i. However, there was no change in the percentage of cells positive for CPSF6 puncta or in the number of puncta per cell over time (Fig S5B).

### The Number of Cells with Puncta Seems Proportional to the Amount of CA That Can Bind CPSF6

As CA could still be detected in non-dividing cells for considerably longer than the peak time of integration, we wondered what oligomeric state the CA was in and how much of the lattice was required to induce puncta formation. We therefore set up a ‘mixed particle’ experiment where we synthesised viral particles by transfecting 293T cells with varying ratios of plasmids expressing WT *gag-pol* or a mutant *gag-pol* carrying the A77V substitution in CA that blocks CPSF6 binding to CA. HeLa cells or MDMs were then infected with these mixed particles for 16h or 72h, respectively, and the number of CPSF6 puncta visualised by immunofluorescence. The number of CA positive HeLa cells with CPSF6 puncta are plotted for each *gag-pol* ratio in Figure S6A. As expected, when 100% of the *gag-pol* contained the A77V CA substitution, the VLPs produced did not induce any CPSF6 puncta upon infection (Figure S6A). The number of puncta increased as more WT *gag-pol* was added to the transfection used to synthesise the VLPs. However, as the number of puncta observed was proportional to the amount of WT *gag-pol* in the transfection, it was hard to determine exactly how much CA is required per particle to form a CPSF6 puncta. A similar result was seen in MDMs (Figure S6B) although the overall number of cells with CPSF6 foci was lower as the infection was likely lower in MDMs and all cells were counted not just CA positive cells (Figure S6B).

### In Macrophages, the Number of Puncta Remain Constant Regardless of the MOI or the Number of Nuclear Speckles

Another way to look at how much CA is required to induce puncta formation is to titrate down the amount of virus used to infect cells. We therefore infected MDMs with different amounts of viral particles and assessed CPSF6 puncta formation by immunofluorescence at different times post infection (Figure 7). Interestingly, we saw that the number of CPSF6 puncta formed per cell remained similar despite varying viral titre and remained similar over the time course of the experiment (Figure 7A-C). The average number of CPSF6 puncta per cell was 2-3 for low MOI infections (Figure 7B) and 2-4 puncta/cell for infections with 40 times more virus (Figure 7C). This is considerably fewer than the average number of CPSF6 puncta seen in HeLa cells (Figure 1). The highest number of puncta seen in any MDM cell in this experiment was eight. This suggests that several viruses may associate with the same CPSF6 puncta in MDMs and that there are limited puncta formed in these cells. However, the number of SC35-containing nuclear speckles does not seem to be the limiting factor as SC35 staining detects many more nuclear speckles in any individual MDM than CPSF6 puncta (Figure 7D).

**Figure 7.**
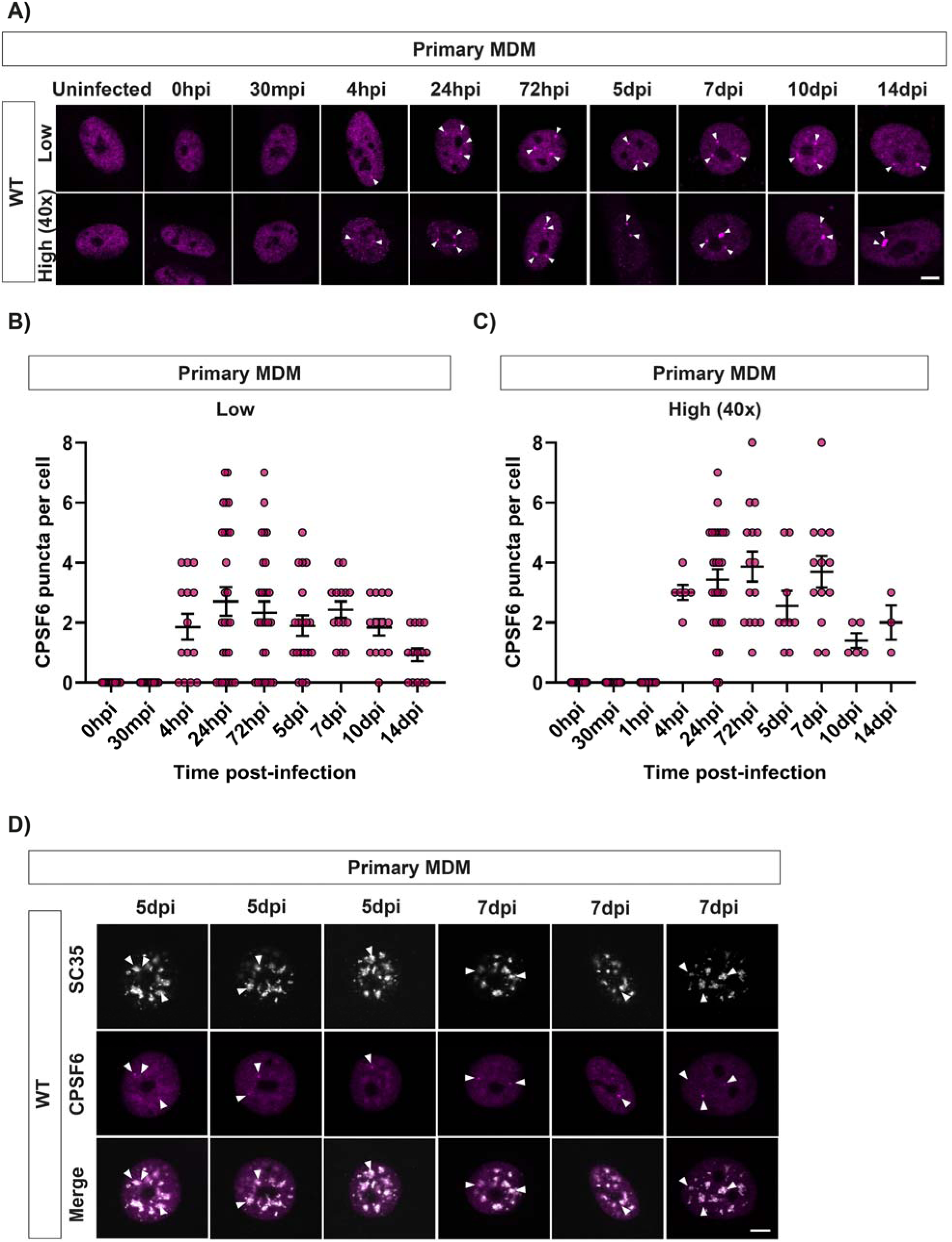
In primary MDM, the number of CPSF6 puncta remain constant regardless of the MOI or the number of nuclear speckles. (A-C) Primary MDMs were synchronously infected with either low (0.3mU RT activity) or high (12mU RT activity) amounts of WT HIV-1-GFP. Cells were fixed at the following times post-infection: 0h, 30min, 4h, 24h, 72h, 5d, 7d, 10d and 14d and immunostained for CPSF6. (A) Representative images are shown. Scale bar is 5μm. (B, C) Graphs show the quantification of the number of CPSF6 puncta per cell in cells infected with low (B) or high (C) RT units of WT VLP. Each dot represents an individual cell. The mean ± SEM from at least two biological repeats is shown. (D) Primary MDMs were synchronously infected with equal RT units of WT HIV-1-GFP and fixed and immunostained for SC35 and CPSF6 at 7 and 10 d.p.i.. Representative images are shown. White arrowheads point to SC35 and CPSF6 co-localisation. Scale bar is 5μm.

## Discussion

Until recently, the core of HIV-1 was thought to uncoat before nuclear entry could occur. Therefore, it was assumed that the CA protein did not participate in nuclear events. However, several groups have now shown that CPSF6, a cellular nuclear protein that binds CA, is re-localised into puncta in the nucleus of infected cells [12,14,18,31,32,37,52]. We previously showed that HIV-1 CA mutants with hyperstable cores get stuck at the nuclear pore and do not re-localise CPSF6, despite being competent to bind CPSF6 in vitro [14]. This implies that at least some CA needs to be present in the nucleus to induce CPSF6 puncta formation, although the oligomeric state of nuclear CA is unknown, evoking the questions: how does CA cause CPSF6 relocalisation and how does this affect HIV-1 replication. Understanding CPSF6 puncta formation better will shed light on the role of CPSF6 in infection.

We have confirmed that CA-dependent CPSF6 relocalisation to SC35-containing nuclear speckles occurs similarly in both dividing and non-dividing cells, including primary MDMs [18,31,37,52]. Interestingly, we observed that CPSF6 already forms puncta that colocalise with SC35 in T-cells, including primary T-cells purified from PBMCs (Figure 2 and Figure S1 and S2). This is noteworthy as T-cells are the major natural target cell of HIV-1 infection. Although the CPSF6 staining did alter slightly, with an increase in smaller, more focused puncta in some instances, we did not see a significant change in the number or distribution of puncta following infection of T-cells with HIV-1 VLPs, suggesting that HIV-1 CA is not driving CPSF6 relocalisation in this cell type. It is not clear why CPSF6 is already punctate in T-cells, but it suggests that HIV promotes and/or mimics a natural behaviour of CPSF6 in other cell types. Interestingly, a recent report also found CPSF6 puncta in several uninfected cell lines [53]. The authors showed that in cancer cells, although overall CPSF6 levels were higher, the CPSF6 underwent less liquid-liquid phase separation (LLPS) resulting in fewer CPSF6 puncta. Reducing the amount of punctate CPSF6 resulted in the preferential usage of proximal poly(A) sites leading to shortened 3’ UTRs of cell-cycle-related genes and accelerated cell proliferation [53]. These results suggest that CPSF6 condensation may modify the function of CPSF6.

Importantly, another component of the CFIm complex, CPSF5, also relocalised along with CPSF6 in both HeLa cells and MDMs (Figure 3 and Figure S3). This is in agreement with other studies [18,52] and suggests that puncta contain other components in addition to CPSF6. It is possible that the whole CPA complex may relocate to nuclear speckles, however, unfortunately, we were unable to determine whether other known CPA proteins relocated to puncta upon infection because either their staining patterns were already grainy/punctate or there was a high level of non-specific background staining. Nevertheless, this has implications for the formation of the CPSF6 biomolecular condensates that have been reported in cells [37,52]. Recent structural studies [38] indicated that low complexity regions in CPSF6 enable avid interactions with CA and mediate CPSF6 multivalent assembly. How CPSF5, and possibly RNA and other proteins in the complex, fit into these assemblies is unknown but it suggests that the viral CA is coated in a considerable amount of cellular protein which could impact core uncoating and nucleic acid release if the shell was still intact at this point.

As another binding partner of CPSF5, CPSF7, did not re-localise with CPSF6 (Figure 3 and Figure S3), this is further evidence that direct binding of CA to CPSF6 drives the formation of these puncta. Interestingly, we detected CA in CPSF6 puncta at very late time points (Figure 6), which for MDMs was several days after integration occurred (Figure 5). This suggests that the CA is stabilised by the CPSF6 puncta and is possibly never released. However, we do not know the oligomeric state of the CA in puncta, and from our mixed particle studies (Figure S6) we can only conclude that puncta formation seems proportional to the amount of wild type CA present. This agrees with a study by Francis et al. [54] who also showed that the efficiency of nuclear transport and integration targeting of HIV-1 was proportional to the amount of WT CA in mixed particles. As low levels of WT CA were sufficient for HIV nuclear accumulation and for targeting integration into speckle-associated genomic domains, this may indicate that partial capsids are sufficient for nuclear events, including CPSF6 puncta formation. Further work will be needed to determine the state of CA in CPSF6 puncta. Furthermore, as we did not image the puncta in live-cell experiments we cannot state whether individual puncta are stable and remain permanently associated with CA, or whether the particles within puncta have integrated. Interestingly, although puncta slowly disappeared in cycling HeLa cells with time (Figure 5), this was linked to cell division as treatment with aphidicolin prevented loss of puncta (Figure 6). An outstanding question is how the presence of CPSF6 puncta in cells for more than two weeks affects the polyadenylation of cellular mRNAs and whether this alters the function of infected MDMs, particularly as altered CPSF6 LLPS affected polyadenylation in cancer cells [53].

In accord with recent reports [37,52], we also showed that CPSF6 puncta form even when reverse transcription and integration are inhibited (Figure 4). Our experiments show that puncta form very rapidly and before the peak of reverse transcription in both HeLa cells and in non-dividing MDMs that have slower reverse transcription kinetics (Figure 5). Scoca et al. have suggested that puncta are required for reverse transcription in macrophages [37]. However, as we have previously demonstrated that mutant viruses with hyperstable cores are able to reverse transcribe [14] but are unable to relocalise CPSF6 (Figure 4), we prefer the notion that reverse transcription happens independently from, although within a similar time frame and location as, puncta formation. The kinetics of reverse transcription likely depend simply on the availability of dNTPs, which would be expected to be more plentiful in the nucleus, and CPSF6 relocalisation appears to be controlled by the presence of CA in the nucleus. Thus, CA seems to drive the trafficking of HIV-1 from the cell periphery right up until the site of integration [30,31,34,35,55] and this occurs without the need for reverse transcription. Although CPSF6 binding directs HIV to integrate into SPADs, future work will determine whether formation of CPSF6 puncta is necessary for integration targeting and how the viral nucleic acid is transferred from CPSF6 puncta to host cell DNA.

An intriguing observation was that there was a limited number of CPSF6 puncta in MDMs (Figure 7). This was not because there was a limited number of nuclear speckles in these cells, as measured by SC35 staining (Figure 7). Nor was it limited by the amount of virus present in the infection as we saw similar numbers of puncta regardless of the viral titre (Figure 7). Prior studies have also shown low numbers of CPSF6 puncta in macrophages [31,32,37], although some have described larger CPSF6 speckles than we noted here. Any differences between studies may be due to differences in fixation protocols for immunofluorescence. We used a combination of PFA and methanol which has been shown to increase CA accessibility during staining [56] whereas other studies only used PFA. Altogether, we propose that in macrophages, CA/CPSF6 is only directed to a fraction of nuclear speckles. Nuclear speckles are known to be dynamic and the 3D spatial organization of genomic DNA around speckles differs between cell types [57]. It will be interesting to establish whether HIV-1 associates with a specific pool of speckles and to determine whether the sites of HIV-1 integration correlate with the DNA associated with these speckles in macrophages.

## Acknowledgments

We thank Melvyn Yap and Jonathan Stoye for helpful discussions and for reading the manuscript.

## Data Availability Statement

The data presented in this study are all contained within the article or supplementary material.

## Author Contributions

Conceptualization, A.G. and K.N.B.; methodology, A.G. and E.R.C.; investigation, A.G., E.R.C. and M.B.; resources, A.G. and E.R.C.; data curation and analysis, A.G., E.R.C., M.B. and K.N.B.; writing—original draft preparation, A.G. and K.N.B; writing—review and editing, A.G., E.R.C., M.B. and K.N.B.; supervision, K.N.B.; project administration, K.N.B.; funding acquisition, K.N.B. All authors have read and agreed to the published version of the manuscript.

## Supplementary Materials

**Figure S1** (associated with Fig 2): CPSF6 staining is punctate in uninfected Jurkat cells and co-localises with nuclear speckles.

**Figure S2** (associated with Fig 2): CPSF6 staining is punctate in uninfected activated primary T-cells and co-localises with nuclear speckles.

**Figure S3** (associated with Fig 3): HIV-1 CA induces relocalisation of CPSF5 and CPSF6 but not CPSF7.

**Figure S4** (associated with Fig 5): CPSF6 puncta form rapidly and decay slowly in HeLa cells.

**Figure S5:** CPSF6 staining is similar in uninfected and infected Jurkat T-cells with time. **Figure S6**: The number of cells with puncta is proportional to the amount of CA that can bind CPSF6.

