## Supplemental file for "HIV-1 Capsid Rapidly Induces Long Lived CPSF6 Puncta in Non-dividing Cells but Similar Puncta Already Exist in Uninfected T-cells"

**Figure S1** (associated with Fig 2)**:** CPSF6 staining is punctate in uninfected Jurkat cells and co-localises with nuclear speckles**.**

**Figure S2** (associated with Fig 2): CPSF6 staining is punctate in uninfected activated primary T-cells and co-localises with nuclear speckles.

**Figure S3** (associated with Fig 3): HIV-1 CA induces relocalisation of CPSF5 and CPSF6 but not CPSF7.

**Figure S4** (associated with Fig 5): CPSF6 puncta form rapidly and decay slowly in HeLa cells.

**Figure S5:** CPSF6 staining is similar in uninfected and infected Jurkat T-cells with time.

**Figure S6**: The number of cells with puncta is proportional to the amount of CA that can bind CPSF6.

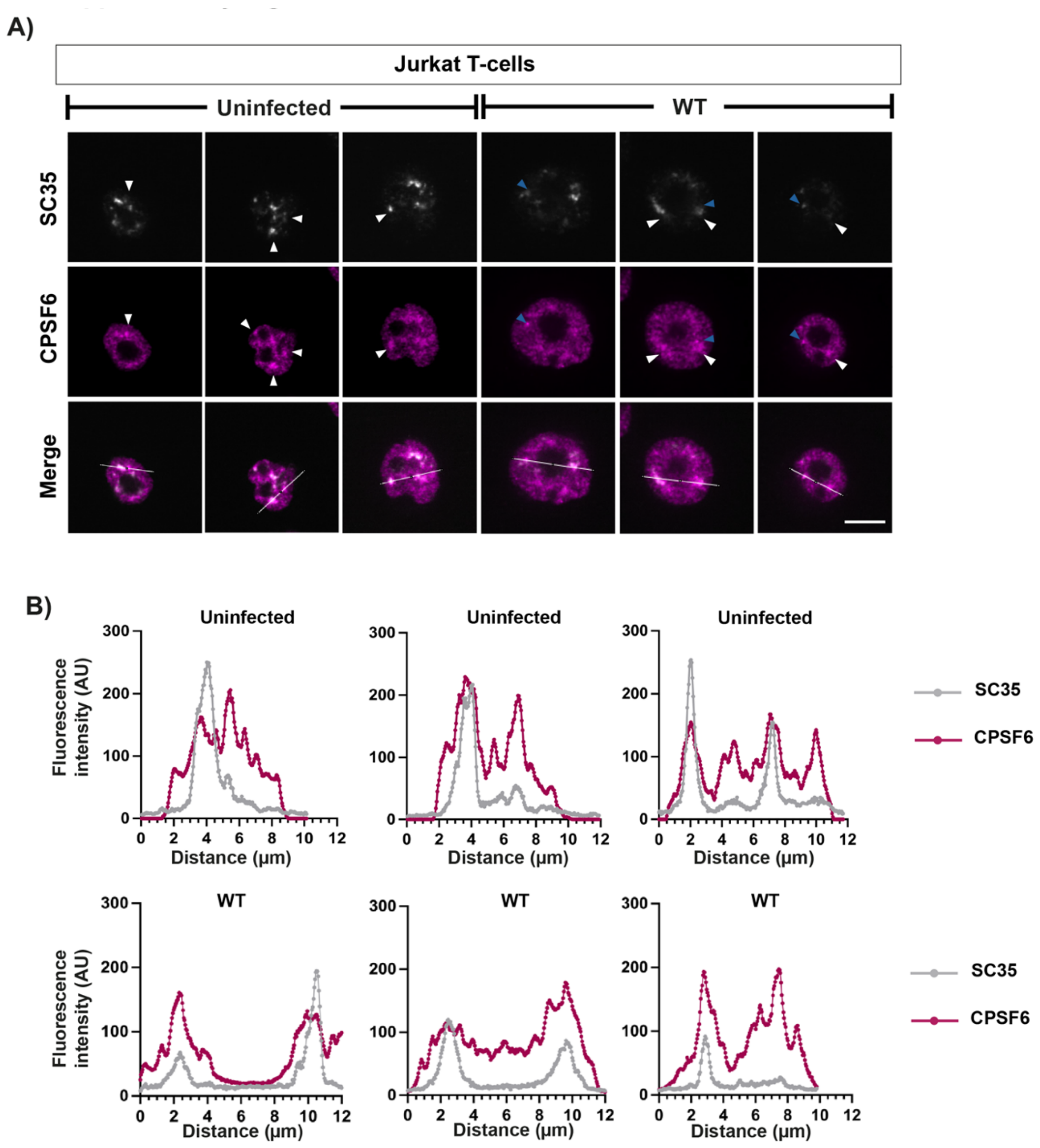


**Figure S1** (associated with Fig 2)**. CPSF6 staining is punctate in uninfected Jurkat cells and co-localises with nuclear speckles.** Jurkat T-cells were infected with WT HIV-1-GFP for 24h, fixed and immunostained for SC35 and CPSF6. (A) Representative images of SC35 (white) and CPSF6 (magenta) staining in uninfected and infected cells. White arrowheads point to SC35 and CPSF6 co-localisation. Blue arrowheads point to small, bright foci. Scale bars are 5 µm. (B) Representative fluorescence intensity profiles of SC35 and CPSF6 staining along the white lines shown in the merged images in (A).

A)

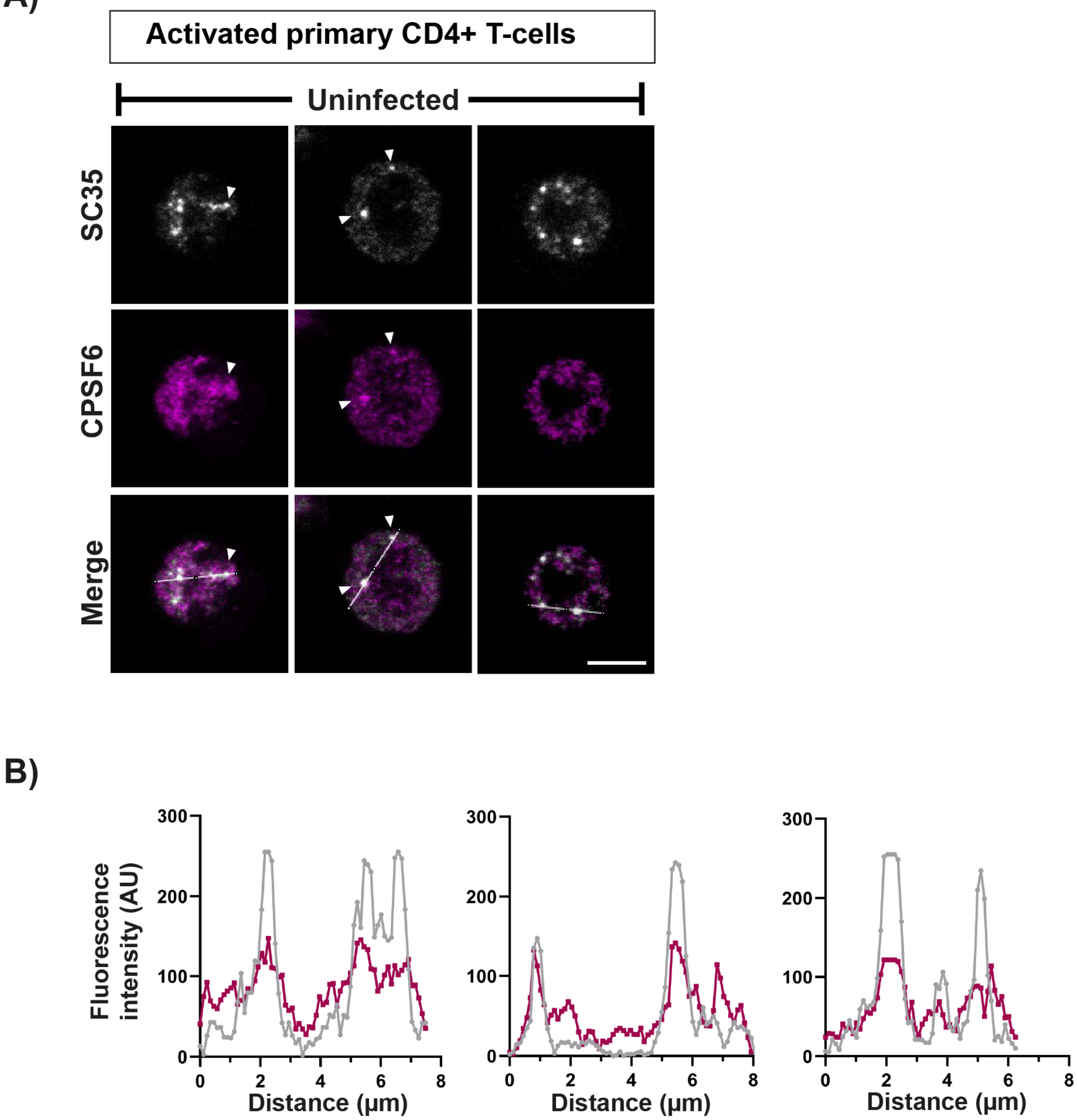


**Figure S2** (associated with Fig 2). **CPSF6 staining is punctate in uninfected activated primary T-cells and co-localises with nuclear speckles.** (A) Primary CD4$^{+}$ T-cells were purified from PBMCs and activated with IL-2 and PHA-L for 72h. After fixation, uninfected cells were immunostained for SC35 and CPSF6. White arrowheads point to SC35 and CPSF6 co-localisation. Scale bars are 5µm.(B) Representative fluorescence intensity profiles of SC35 and CPSF6 staining along the white lines shown in the merged images in (A).

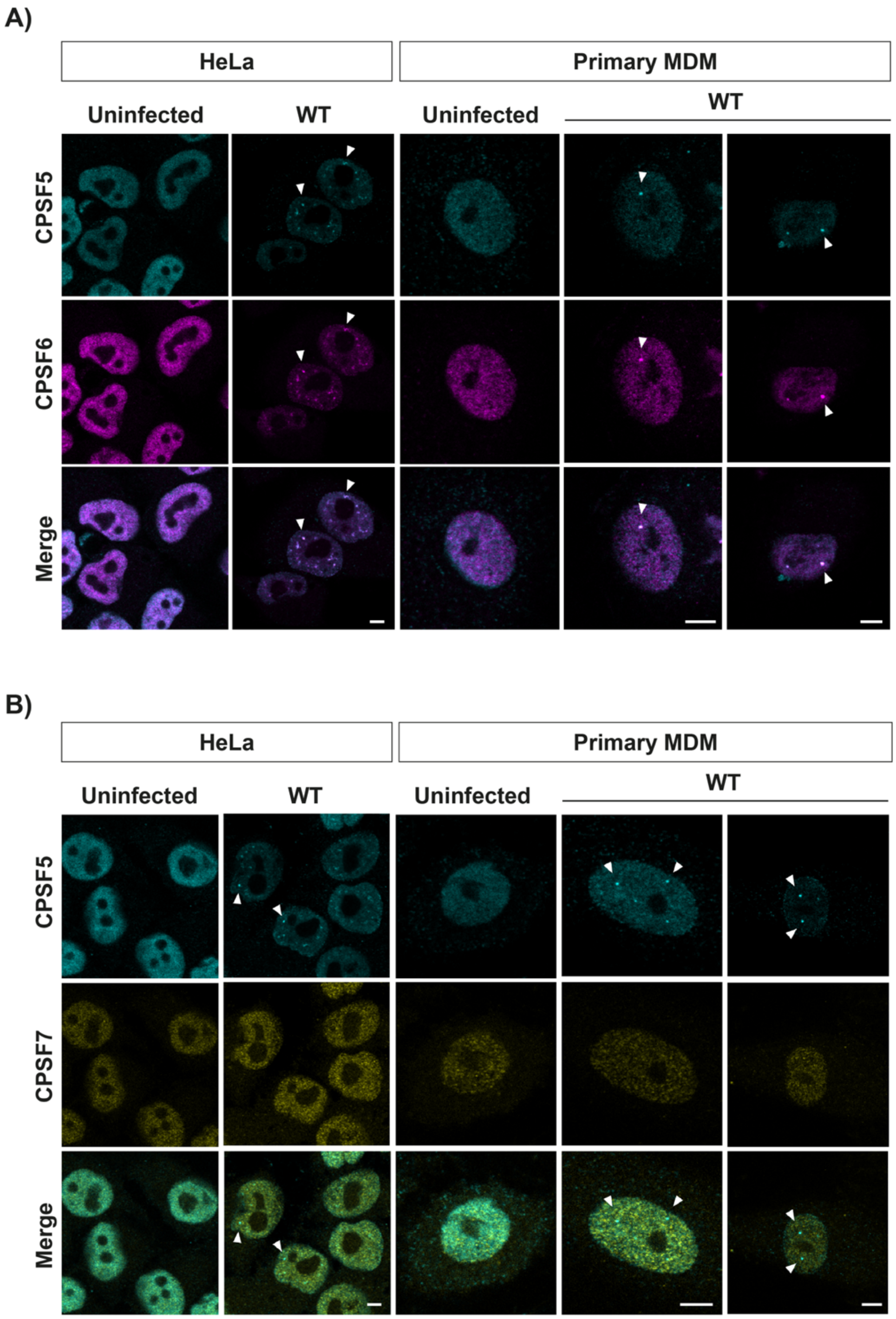


**Figure S3** (associated with Fig 3)**. HIV-1 CA induces relocalisation of CPSF5 and CPSF6 but not CPSF7.** HeLa cells and primary MDMs were infected with equal RT units of WT HIV-1-GFP and fixed at 16 h.p.i. and 72 h.p.i., respectively. Uninfected and infected cells were immunostained for CPSF5 and CPSF6 (A) or CPSF5 and CPSF7 (B). Representative images are shown. White arrowheads point to co-localisation. Scale bars are 5µm.

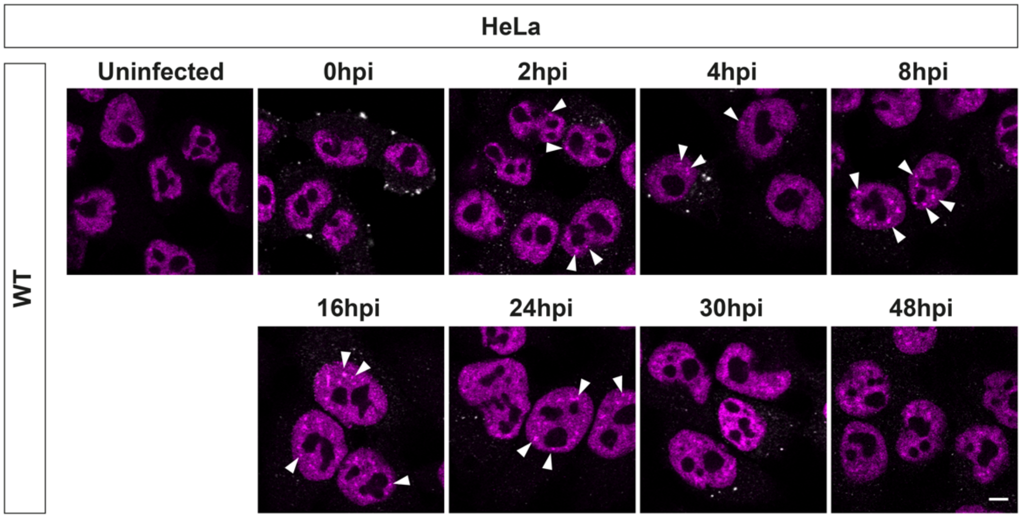


**Figure S4** (associated with Fig 5). **CPSF6 puncta form rapidly and decay slowly in HeLa cells.** HeLa cells were synchronously infected with equal RT units of WT HIV-1-GFP, and fixed at 0, 2, 4, 8, 16, 24, 30 and 48 h.p.i. Cells were immunostained for HIV-1 CA (white) and CPSF6 (magenta). Representative images corresponding to data in Fig 5A are shown. Scale bar is 5µm.

A)

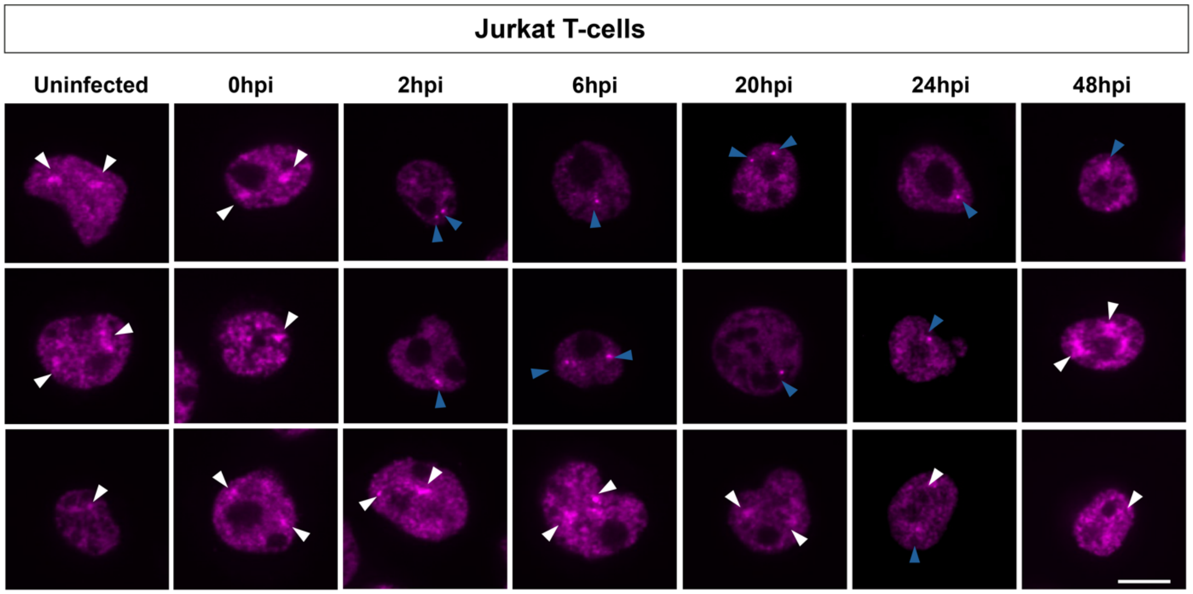


B)

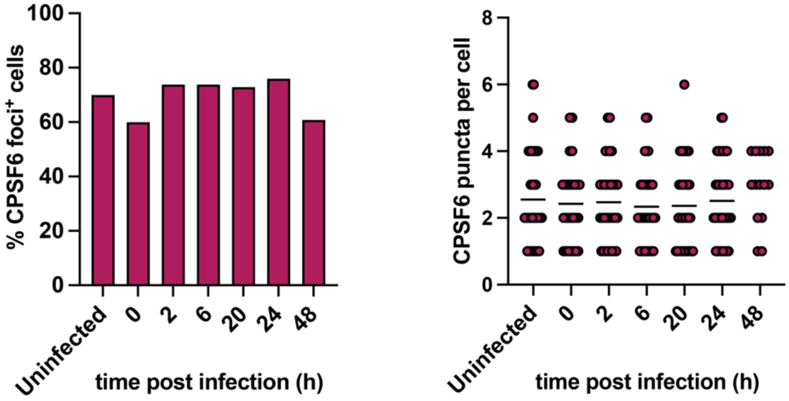


**Figure S5. CPSF6 staining is similar in uninfected and infected Jurkat T-cells with time**. Jurkat T-cells were synchronously infected with equal RT units of WT HIV-1-GFP and fixed at 0, 2, 6, 20, 24 and 48 h.p.i. (A) Representative images of CPSF6 staining (magenta). White arrowheads point to CPSF6 puncta. Blue arrowheads point to small, bright foci. Scale bars are 5 µm. (B) Quantification of the percentage of cells positive for CPSF6 foci (left graph) and the number of CPSF6 puncta per cell (right graph).

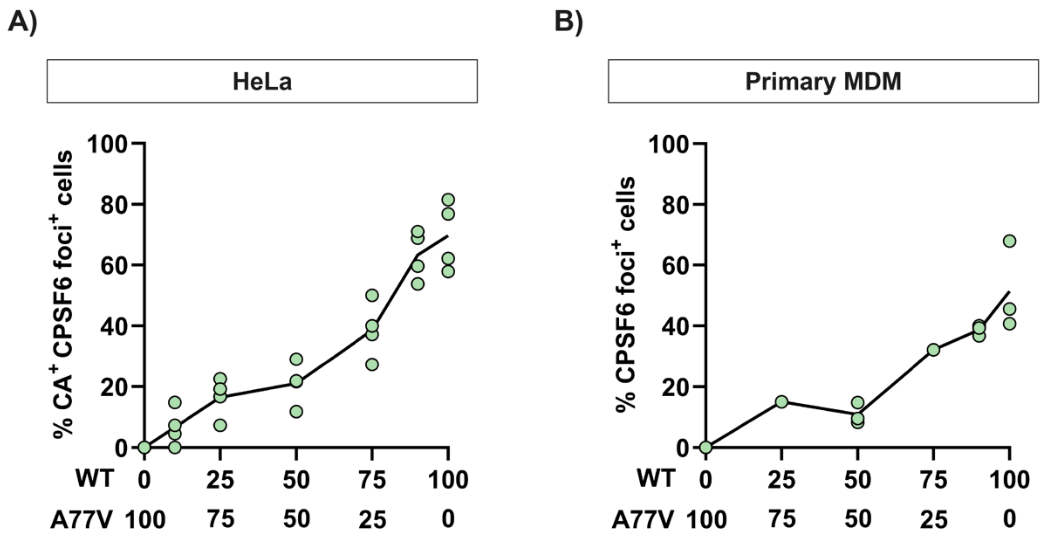


**Figure S6. The number of cells with puncta is proportional to the amount of CA that can bind CPSF6.** HeLa (A) or primary MDMs (B) were synchronously infected with equal RT units of WT, A77V or "mixed" HIV-1-GFP virions for 16h or 72h, respectively. "Mixed" virions were produced by transfecting 293T cells with varying ratios of plasmids expressing WT *gag-pol* or mutant CA-A77V *gag-pol* as indicated on the x-axis. Cells were immunostained for HIV-1 CA and CPSF6. Graphs show the percentage of (A) CA positive cells or (B) total cells that showed CPSF6 redistribution to puncta. Individual points represent different biological repeats (n=4 for HeLa and n=3 for primary MDM) and the line shows the mean.
